# Heterotypic competition between cancer cells and hepatocytes generates heterogeneous context-dependent phenotypes

**DOI:** 10.1101/2024.05.15.593526

**Authors:** Andrea Spinazzola, Tânia Carvalho, Miguel Alexandre Ferreira Pinto, Mariana Marques-Reis, Andrés Gutiérrez-García, Davide Accardi, Eduardo Moreno

## Abstract

Competitive interactions between tumor cells and surrounding healthy cells are constantly present during the progression of a solid tumor, and their outcome has been proposed to affect the clinical behavior. Previous studies have described various mechanistic and molecular aspects that characterize this process, overall indicating that cancer cells behave as supercompetitors, which eliminate neighboring healthy cells to gain vital space for growth and infiltration of the tissue. Nevertheless, there is a lack of systematic characterization of these competitive interactions, particularly in the context of cancer in mammals. Furthermore, previous studies in the field of cell competition have primarily focused on homotypic cell competition, involving different clones of the same cell or cells deriving from the same tissue. Data are scarce regarding heterotypic cell competition between two unrelated cell types, which is particularly critical for the understanding of metastatic tumors. In this research, we study cell competition in the context of liver metastases, providing a broad characterization of this process in different relevant scenarios, including cells growing *in vitro* in 2D and 3D, and *in vivo*. Results show that *in vitro*, only a subset of cancer cell lines are coherently strong or moderate competitors against hepatocytes, while the remaining demonstrate poor competitiveness. The competitive proficiency can vary depending on the experimental growth system that is employed, and often predicts the phenotype of liver metastases in terms of aggressiveness and morphology. Finally, our data point towards an involvement of mechanical competition in determining the supercompetitor trait of cancer cells. Altogether, our research provides the first comprehensive characterization of heterotypic cell competition, and indicates that cancer cells possess heterogeneous competitive proficiency towards hepatocytes which can be affected by the growth conditions.

## Introduction

The destruction of the host tissues and their replacement with tumor is a common morphological finding in various types of advanced solid cancers, including both locally advanced primary tumors and metastases [1–4]. To explain this phenomenon, it has been proposed that cancer cells behave as supercompetitors, eliminating and replacing surrounding healthy cells through a process called cell competition [5–12]. Cell competition is a widespread phenomenon, conserved across species, in which cells sense their fitness status, and the confrontation of cells with different fitness levels drives the elimination of those cells (named as loser) that are less fit than their neighbors (named as winner).

Competitive interactions between transformed cells and the surrounding host cells are constantly present during all phases of cancer development and progression [5,13,14]. The outcome of these interactions has been suggested to be relevant in determining various aspects of the disease, including the tumor morphological features, the clinical behavior, and the response to anticancer therapy. Although cell competition generally acts as a tumor-suppressive mechanism during the early stages of carcinogenesis [27,28,31–34], cancer cells can hijack this process to promote their own growth and dissemination throughout the organism by becoming supercompetitors [5–10,13]. Available evidence suggests that supercompetitor tumor cells can exploit different types of cell competition to eliminate and replace neighboring healthy cells and infiltrate host tissues [14,15]. These include flower (Fwe) fitness fingerprint-mediated cell competition [10,11], tumor expression of death ligands, in particular Fas ligand (FasL) [16–21], and cannibalism of the host cells [22]. Moreover, mechanical cell competition is predicted to have a strong impact on the initiation and the progression of cancer [23,26], although direct evidence in mammalian tumors is still lacking.

An important limitation of the published data is the absence of a broad phenotypic and mechanistic characterization of cell competition in mammalian tumors. In fact, previous studies evaluated only one or a few tumor models, yet most of them have been conducted focusing only on one single mechanism of cell competition. Importantly, most knowledge in the field derives from the study of cell competition between different clones of the same cell type or cells deriving from the same tissue. We define this setting as homotypic cell competition. However, to understand how cell competition affects the pathophysiology of metastatic tumors, it is crucial to study this process in the context of two unrelated cell types, namely heterotypic cell competition.

In this work, we decided to address all these issues by studying the competitive interaction between cancer cell lines, originating from different tissues, and hepatocytes, which is relevant in the context of liver metastasis. The goal is to provide a broad characterization of the process, and explore the molecular mechanisms that lead to the elimination of the hepatocytes.

## Results

### Heterotypic cell competition in 2D culture

To systematically study the competitive interactions between cancer cells and hepatocytes, we set up a cell competition assay *in vitro*. We started by culturing together, as 2 dimensional (2D) monolayers, the murine hepatocyte cell line AML12 expressing EGFP and cancer cell lines expressing TdTomato (TdT). Cells were seeded subconfluent, at a ratio AML12 cell-cancer cell of 3:1 for murine tumor cells and 2:1 for human tumor cells. The culture was periodically monitored for 12 days through imaging with a confocal microscope. As a readout for the population size of AML12 cells during the experiment, we measured the total area per field of view occupied by GFP-labeled cells and quantified its variation.

We assessed eight murine cancer cell lines, representing colorectal carcinoma (MC38 cells and CT26 cells), pancreatic adenocarcinoma (Pan02 cells), melanoma (B16 cells), mammary carcinoma (4T1 cells and EO771 cells), lung cancer (LLC cells) and renal cancer (Renca cells), which are tumors that typically metastasize to the liver. We found that only two of them, 4T1 cells and B16 cells, behave as strong competitors, resulting in a substantial loss (>60%) of AML12^EGFP^ cells from the culture of approximately 85% and 97%, respectively [Fig. 1A-B and fig. S1A]. In parallel, B16 cells and 4T1 cells colonized the whole culture area within a few days. On the other hand, MC38 cells, Pan02 cells, LLC cells, and CT26 cells lead only to a minimal decrease (≤20%), or even to an increase in the AML12^EGFP^ cells population, and colonized only a fraction of the culture area. Accordingly, we define these cell lines as “poor competitors”, in the sense that they coexist with AML12 cells without major competitive outcomes. EO771 cells showed a moderate competitive proficiency, with a slow but consistent loss of AML12 cells and the colonization of the majority of the culture area. Finally, Renca cells induced a significant loss of AML12^EGFP^ cells and colonized around half of the culture area during the first 8 days. This was followed by a stall, without major changes in both cell populations, which lasted until day 20 of culture, when we ended the experiment [Fig. S1A and data not shown].

**Figure 1:**
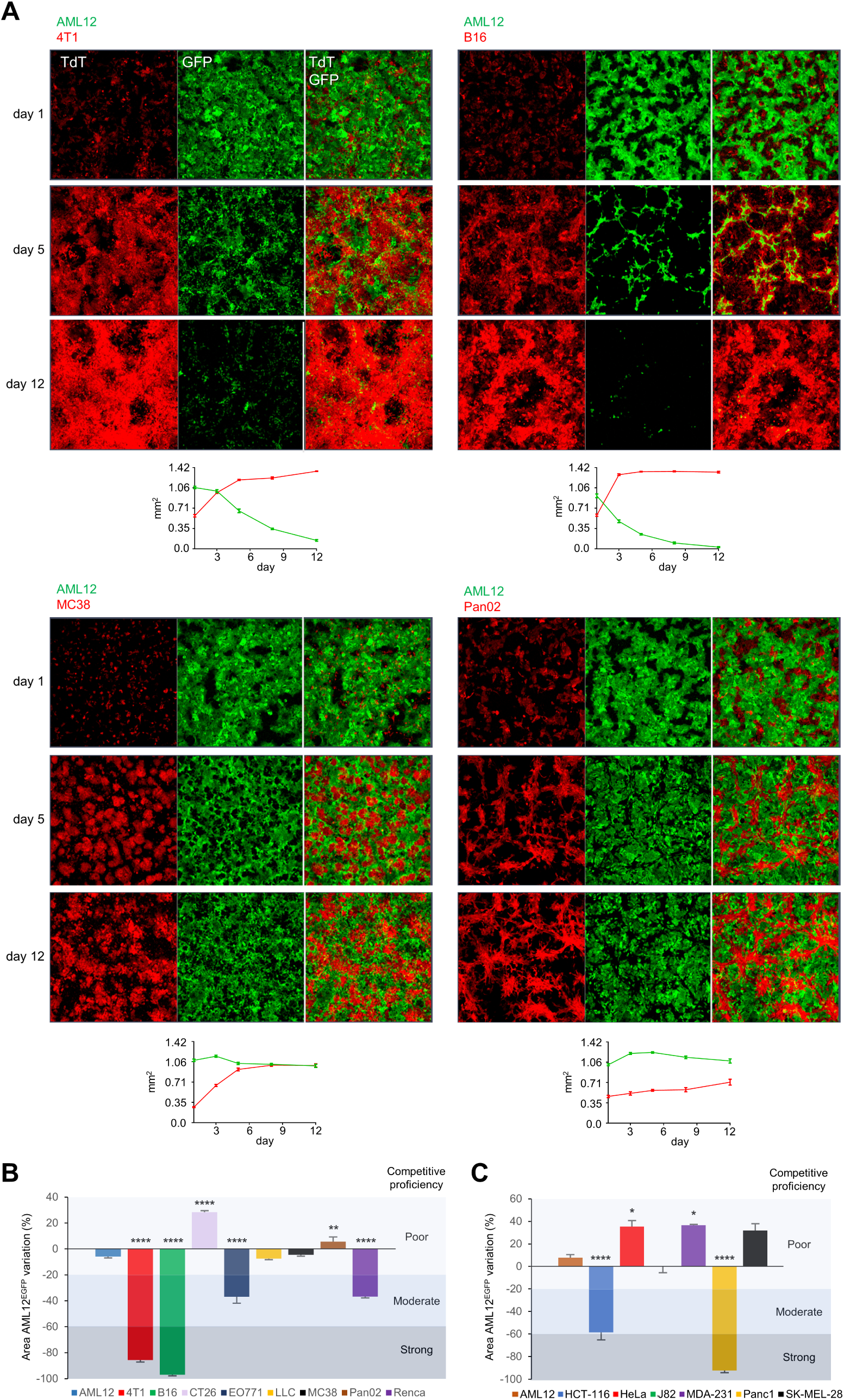
Competition between cancer cells and AML12 cells in 2D culture. **(A)** Representative images, shown as maximum intensity projection, of AML12^EGFP^ cells (green) in culture with murine cancer cells expressing TdTomato (TdT, red). At the bottom of each panel, a chart shows the area per field of view (10x objective) occupied over time by AML12^EGFP^ cells (green curve) and the partner cell line (red curve). Cells were seeded in duplicate or more, each dot shows the mean and the SEM of the values from at least 4 photos. **(B)** Variation of the total area per field of view occupied by AML12^EGFP^ cells in culture with murine cancer cells, measured as the ratio 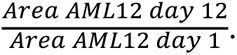 Each color corresponds to a different partner of culture of AML12^EGFP^ cells, as indicated at the bottom of the graph. The variation obtained with each cancer cell line is compared with the one measured with AML12^TdT^ cells. Cancer cells are defined as poor competitors when the loss of AML12^EGFP^ cells is ≤20%, strong competitors when the loss is >60%, and moderate competitors when the loss is between 21% and 60%. p-values for ANOVA test are shown with the following designation: *p<0.05, **p<0.01, ***p<0.001, ****p<0.0001. **(C)** Variation of the area per field of view occupied by AML12^EGFP^ cells in culture with human cancer cells.

When we cultured six human cancer cell lines with AML12 cells, only pancreatic adenocarcinoma Panc1 cells behaved as strong competitors, eliminating approximately 92% of AML12 cells. Colorectal cancer HCT-116 cells resulted moderate competitors, while cervical cancer HeLa cells, urothelial carcinoma J82 cells, breast cancer MDA-231 cells and melanoma SK-MEL-28 cells were poor competitors [Fig. 1C and fig. S2].

We next tested whether cancer cells need to physically interact with AML12 cells to outcompete them. First, we cultured AML12 cells alone and treated them with the conditioned media derived from cancer cells. This had no detrimental effect on AML12 cells population, regardless of the cancer cell line that produced the conditioned media [Fig. S1B]. We then cultured AML12 cells on top of a permeable transwell insert which was placed inside a multiwell plate containing cancer cells growing at the bottom. In this way, AML12 cells and cancer cells are cultured together and share the same media, however they are not in contact. Over 8 days of culture, strong competitor B16 cells and 4T1 cells did not decrease the size of the population of AML12 cells [Fig. 2A]. To exclude the presence of soluble factors that are produced by cancer cells specifically when they are in contact with AML12 cells, we treated AML12 cells with the supernatant collected from cocultures with cancer cells. Again, their population size was not reduced by the addition of the supernatant from cocultures with 4T1 cells and B16 cells [Fig. S1C]. Taken together, these data indicate that the competition between cancer cells and AML12 cells is contact-dependent and is not executed through paracrine signaling.

**Figure 2:**
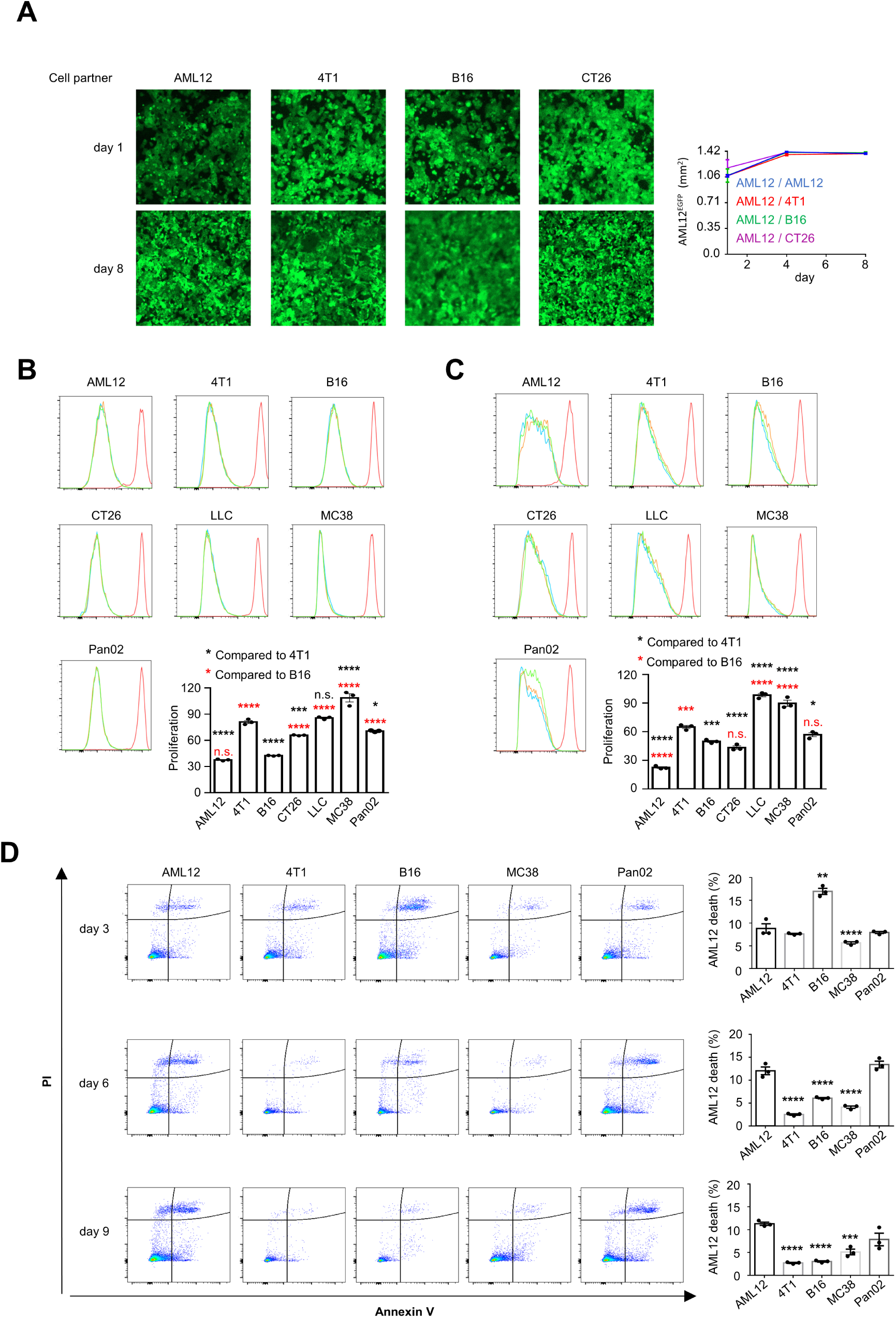
Phenotypic characterization of heterotypic cell competition in 2D culture. **(A)** On the left, representative images of AML12^EGFP^ cells cultured on top of a permeable insert placed inside a multiwell plate containing the indicated cell lines growing at the bottom. On the right, chart of the total area per field of view occupied by AML12^EGFP^ cells over 8 days of culture. **(B)** Flow cytometry histograms and quantification of the proliferation of murine cells in monoculture, assessed as the mean dilution of cell trace violet (CTV) at 72 hours after seeding. The experiment was done in triplicate. p-values are obtained with ANOVA test. **(C)** Flow cytometry histograms and quantification of the proliferation of murine cells in culture with AML12^EGFP^ cells for 96 hours. **(D)** Representative flow cytometry dot plots and quantification of AML12^EGFP^ cells undergoing cell death after 3 days, 6 days and 9 days in coculture with cancer cells as indicated on the top. The total amount of AML12 cells death is obtained through the sum of the percentage of AML12^EGFP^ cells that are positive for Annexin V, propidium iodide (PI), or both. The percentage of dead AML12^EGFP^ cells in culture with each cancer cell line is compared with that obtained with control AML^TdT^ cells. p-values are obtained with ANOVA.

To investigate whether the different phenotypes that we observed were a consequence of the proliferation rate of cancer cells, we marked tumor cells with Cell Trace Violet (CTV), and quantified by flow cytometry the dye dilution after 3 days of growth as monoculture, and 4 days of growth in culture with subconfluent AML12 cells. Results show that the strong competitors are not the most proliferative cell lines [Fig. 2B-C]. In fact, both 4T1 cells and B16 cells generally proliferated slower than, or at the same rate as poor competitor cancer cells, both as monoculture and in culture with AML12 cells. Similarly, when we quantified the total mass of cancer cells monocultures through crystal violet staining, B16 cells had a lower value than the poor competitors LLC cells, MC38 cells and Pan02 cells [Fig. S1D]. These data argue that the proliferation rate of cancer cells is not sufficient to explain their supercompetitor behavior.

At last, we sought to determine the amount of AML12^EGFP^ cells undergoing apoptosis by flow cytometry. After three, six, and nine days of coculture, we collected the cells and stained them with Annexin V and propidium iodide (PI). To our surprise, we could not detect increased apoptosis of AML12^EGFP^ cells in culture with any cancer cell line compared to control AML12^TdT^ cells, with the exception of B16 cells at day 3 of culture [Fig. 2D]. This suggests that processes different from apoptosis might mediate the elimination of AML12 cells in this context, for example apical extrusion or other forms of cell death which are not detected by the assay that we used.

### Heterotypic cell competition in 3D culture

Culturing cells in 2D has a multitude of limitations concerning its inability to emulate *in vivo* conditions and provide physiological relevance [25]. Accordingly, we decided to further study the competitive phenotype of cancer cells in a 3 dimensional (3D) culture, by growing cell spheroids on top of an extracellular matrix. To monitor the changes in the population size of AML12^EGFP^ cells, we quantified the total volume occupied by GFP per field of view. We observed that AML12 cells stop proliferating shortly after forming spheroids, while all the cancer cell lines sustain proliferation. After 12 days of culture, B16 cells and 4T1 cells maintained their state of strong competitors, resulting in an extensive (>75%) loss of AML12^EGFP^ cells of about 100% and 83%, respectively [Fig. 3A-B]. Similarly, CT26 cells, MC38 cells, and Pan02 cells maintained their status of poor competitors, leading to a loss of AML12^EGFP^ cells ≤50% [Fig. 3A-B and fig. S3]. We set this threshold as approximately twice the loss resulting from the culture with AML12^TdT^ cells (about 27%). EO771 cells also confirmed their status as moderate competitors, generating a loss of about 68%. Strikingly, although LLC cells were poor competitors when cultured in 2D, they exhibited a shift to moderate competitiveness in 3D, eliminating about 64% of AML12^EGFP^ cells. Conversely, Renca cells were intermediate competitors in 2D but transitioned to poor competitiveness in 3D culture [Fig. 3B and fig S3].

**Figure 3:**
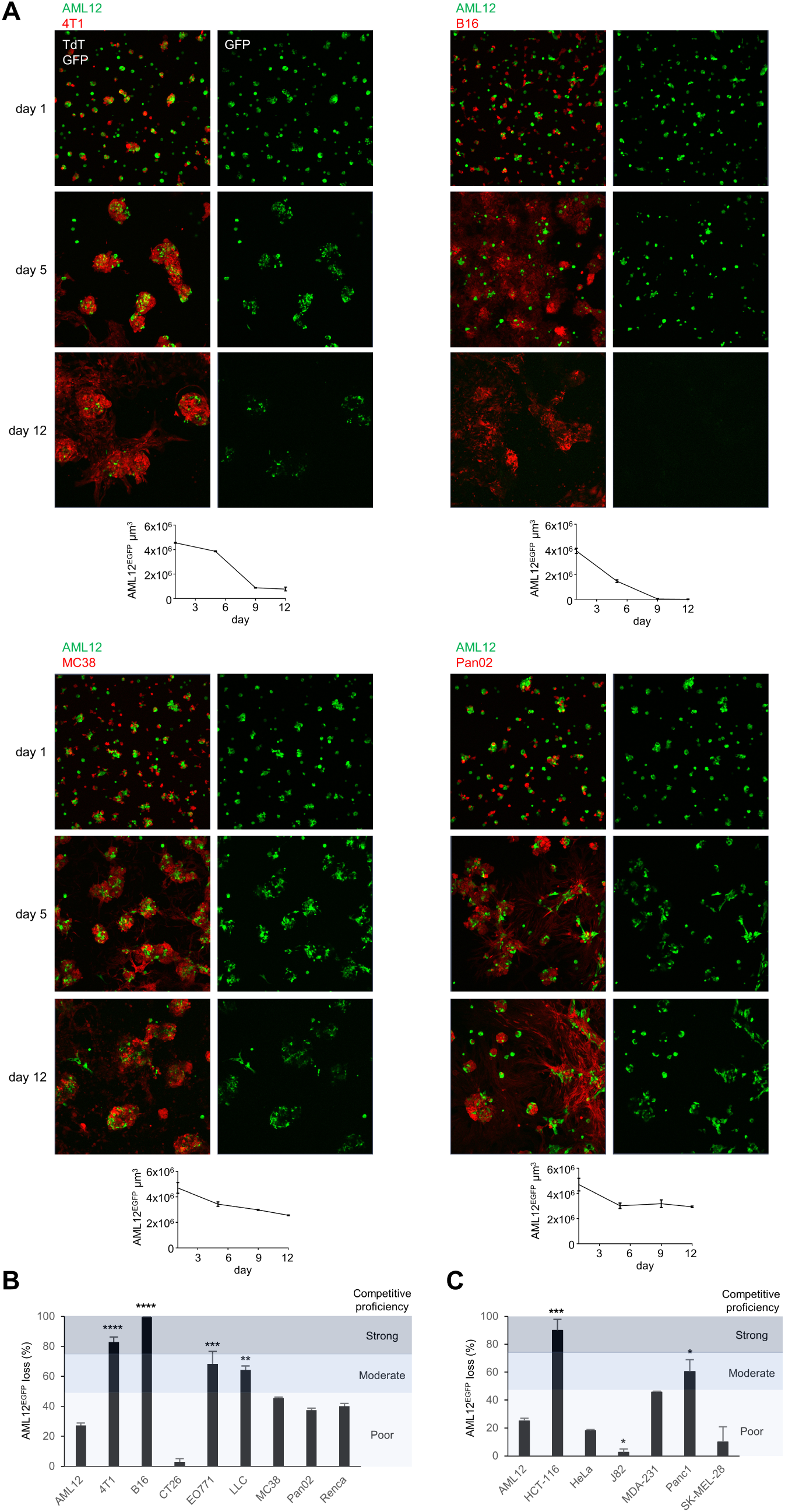
Competition between cancer cells and AML12 cells in 3D culture. **(A)** Representative images, shown as maximum intensity projection, of AML12^EGFP^ cells (green) in culture with murine cancer cells marked with TdTomato (TdT, red) on top of Matrigel gel. At the bottom of each panel, a chart shows the total volume per field of view (10x objective) occupied over time by AML12^EGFP^ cells. Each dot represents the mean and the SEM from at least 3 images. **(B)** Quantification of the loss of AML12^EGFP^ cells in 3D culture with murine cancer cells, measured as the ratio 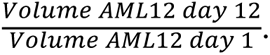 At the bottom of each column, the partner cell in culture with AML12^EGFP^ cells is listed. The loss with each cancer cell line is compared to that obtained with AML12^TdT^ cells. Cancer cells are categorized as strong competitors when the loss of AML12^EGFP^ cells is >75%, while they are considered poor competitors when the loss is ≤50%. p-values for ANOVA test are shown. (**C)** Quantification of the loss of AML12^EGFP^ cells in 3D culture with human cancer cells.

To confirm that the elimination of AML12 cells was an active process and was not due to detachment from the top of the extracellular matrix, we tested a fully embedded coculture of AML12^EGFP^ cells with cancer cells. In agreement with the previous experiment, both 4T1 cells and B16 cells induced a substantial loss of AML12 cells of about 70% and 77%, respectively, which was approximately twice as high as the loss observed with the poor competitors MC38 cells and Pan02 cells [Fig. S4A-B].

Interestingly, all cancer cell lines formed spheroids which consistently incorporated AML12 cells [Fig. 3A and fig. S3]. To dynamically investigate this interaction, we conducted time-lapse microscopy for 42 hours, beginning 3 days after seeding [Supplementary video 1]. Spheroids composed by AML12^EGFP^ cells and AML12^TdT^ cells were predominantly static in terms of movement and morphology, and occasionally exhibited fragmentation of AML12^EGFP^ cells, a sign of cell death. In contrast, spheroids containing cancer cells typically displayed intense movements and shape modifications. Remarkably, they appeared to actively attract and incorporate AML12^EGFP^ cells, both as single cells and as spheroids, especially in cultures with 4T1 cells, CT26 cells and MC38 cells. Fragmentation of AML12^EGFP^ cells was often detected in the culture with B16 cells, while it was infrequent with the other cancer cell lines.

Among human cancer cells, only HCT-116 cells behaved as strong competitors, leading to a near complete elimination (approximately 90%) of AML12 cells [Fig. 3C and fig. S5]. Panc1 cells shifted to a moderate competitor phenotype, while HeLa cells, J82 cells, MDA-231 cells, and SK-MEL-28 cells all maintained their status of poor competitors.

Similar to what we observed in 2D culture, when we treated AML12 cells spheroids with the conditioned media derived from cancer cells there was no effect on their population size [Fig. S4C]. The same happened when we treated AML12 cells spheroids with the supernatant collected from a coculture of cancer cells with AML12 cells [Fig. S4D]. These data further confirm that the competition between cancer cells and AML12 cells is contact-dependent.

Taken together, these experiments show that cancer cells display a heterogeneous degree of supercompetition towards hepatocytes, which is contact-dependent and can vary depending on the culture system that is employed.

### Strong competitor cancer cells generate aggressive liver metastases with infiltrative morphology

Experimental liver metastases were induced by injecting murine cancer cells into the spleen of syngeneic adult mice. Animals were sacrificed 21 days after the injection of tumor cells, or earlier upon reaching humane endpoints. All the cell lines generated macroscopic liver metastases in 100% of the mice, with generally low intra-group variability in terms of tumor burden and clinical course. Notably, the three cell lines with a consistent supercompetitor trait *in vitro*, which was coherent in 2D culture and 3D culture, were also the ones with the most aggressive phenotype *in vivo*, resulting in early termination of the experiment due to reaching humane endpoints at day 12 for B16 cells and EO771 cells, and at day 14 for 4T1 cells. In contrast, none of the mice injected with LLC cells, MC38 cells, Pan02 cells showed clinical signs of disease, except for progressive hepatomegaly, and all were sacrificed in the experimental endpoint, or on day 20 for Renca cells [Fig. 4A-C].

**Figure 4:**
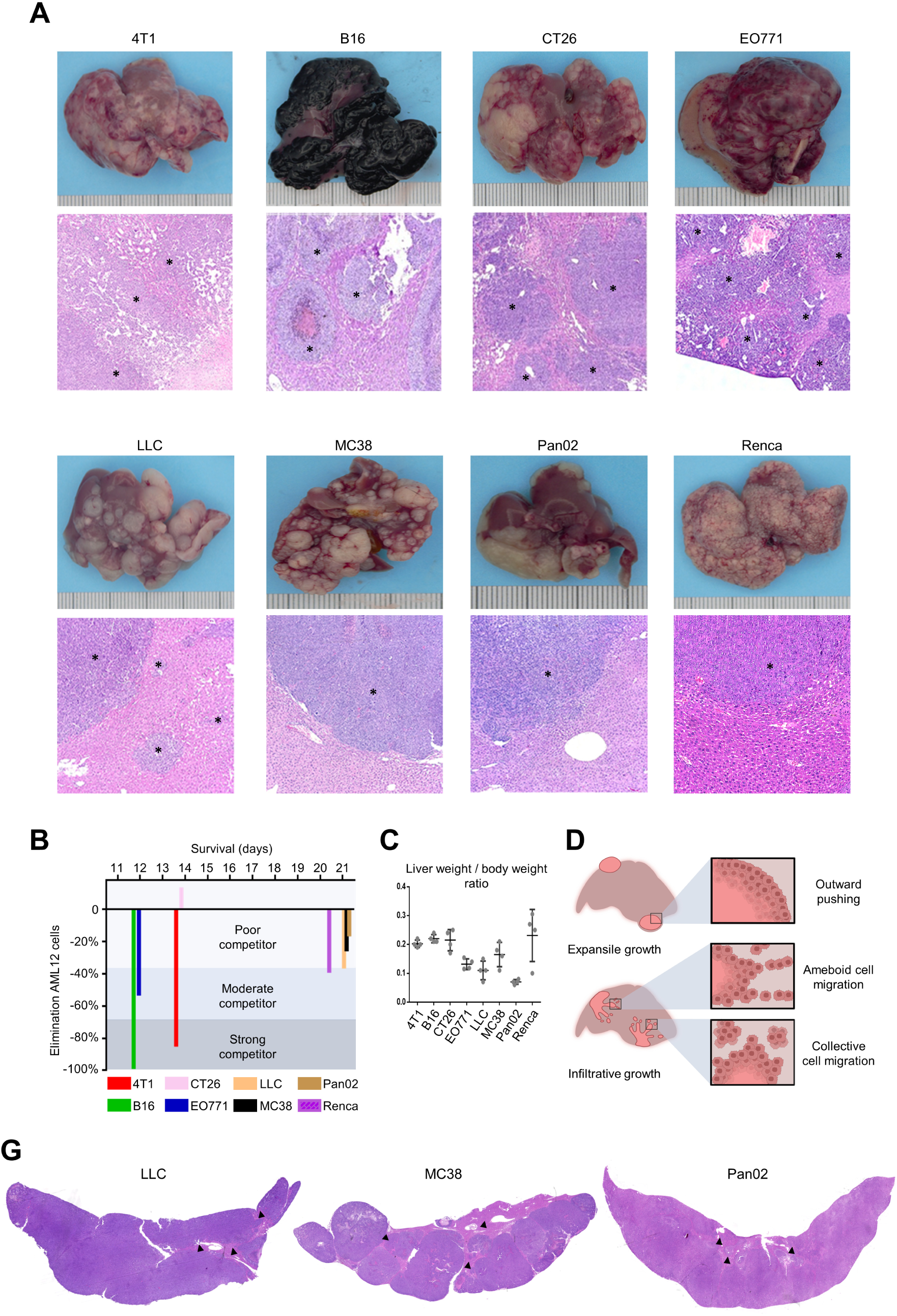
Experimental liver metastases. **(A)** Representative images of liver specimens and corresponding sections stained with hematoxylin and eosin. Tumor lesions are indicated with an asterisk. **(B)** Graph plotting together the sum of the variation of AML12^EGFP^ cells population in 2D culture and 3D culture with the survival of the mice injected with the respective cancer cell line. **(C)** The tumor load is quantified as the ratio between the weight of the liver and the weight of the mouse on the day of sacrifice. The chart shows the mean and the SEM of 4 mice per tumor type. **(D)** Schematic illustration of the morphological characteristics of liver metastases. Image created with BioRender.com. **(G)** Representative images of liver sections vastly replaced by metastatic tumor cells (areas with residual hepatocytes are indicated with an arrowhead).

We found high heterogeneity in the morphological features of liver lesions from different cancer types. There was, on the other hand, high homogeneity in the features seen in the different animals from the same experimental group. Histopathological parameters like tumor growth pattern, tumor border configuration, intercalation between cancer cells and hepatocytes, and the presence of liver damage were investigated and are resumed in Table 1 and fig. 4D. Metastases deriving from the strong competitor 4T1 cells and B16 cells grew endophytically and were highly infiltrative, with cancer cells migrating in small groups inside the liver. These tumors markedly intercalated with the hepatocytes, and associated with marked single cell necrosis of the entrapped hepatocytes. In contrast, the poor competitor MC38 and Pan02 models are characterized by multifocal expansile or poorly infiltrative tumors, respectively, which are well demarcated, with little to no infiltration. Intercalation with the hepatocytes was minimal to moderate, and liver damage was minimal. Remarkably, although CT26 cells were poor competitors *in vitro*, they generated aggressive metastases that required the sacrifice of mice at day 14 due to massive hepatomegaly, and were morphologically similar to those from 4T1 cells and B16 cells. This again suggests that the competitive behavior of cancer cells is context-dependent, potentially influenced by the complex *in vivo* microenvironment, and can differ depending on whether cells grow *in vitro* or *in vivo*. In support of this hypothesis, we also observed instances of liver sections vastly replaced by cancer cells in metastases derived from poor competitor MC38 cells and Pan02 cells, and from poor-moderate competitor LLC cells [Fig. 4G], indicating that cancer cells that do not proficiently eliminate hepatocytes *in vitro* may do it *in vivo*.

**Table 1.** Histopathological features of experimental liver metastases.

|  | 4T1 | B16 | CT26 | EO771 | LLC | MC38 | Pan02 | Renca |
| --- | --- | --- | --- | --- | --- | --- | --- | --- |
| Histology | Breast cancer (high grade) | Melanoma | Colorectal cancer (high grade) | Breast cancer (intermediate grade) | Lung cancer | Colorectal cancer (low grade) | Pancreatic cancer | Renal cancer |
| Growth pattern | Infiltrative | Infiltrative | Infiltrative | Infiltrative | Infiltrative | Expansile | Mixed | Infiltrative |
| Tumor-liver interface | Collective cancer cell migration | Collective cancer cell migration | Collective cancer cell migration | Ameboid cancer cell migration | Collective and ameboid cancer cell migration | Outward pushing | Outward pushing and collective cancer cell migration | Outward pushing and collective cancer cell migration |
| Intercalation | Marked | Marked | Marked | Marked | Marked | Mild | Moderate | Mild |
| Liver damage | Marked single cell necrosis of entrapped hepatocytes | Marked single cell necrosis of entrapped hepatocytes | Marked hepatocellular vacuolation, hyperplasia and inflammation | Mild single cell necrosis of entrapped hepatocytes | Moderate compressive/ischemic necrosis, mild single cell necrosis of entrapped hepatocytes | Moderate compressive/ischemic necrosis | Mild hepatocellular hyperplasia | Minimal single cell necrosis of entrapped hepatocytes |

Immunostaining of liver sections with anti-cleaved Caspase-3, to detect apoptosis, showed only rare positive hepatocytes inside the tumor and at the tumor margin, at a similar number as in liver parenchyma remote from the tumor [Fig. S6]. It is therefore possible that the single cell necrosis seen in entrapped hepatocytes within tumors occurs via a caspase-independent form of cell death. At last, we also attempted to generate xenograft liver metastases by injecting human cancer cells into the spleen of nude mice. However, out of 5 cell lines tested, metastases were only detected in one of the mice injected with MDA-231 cells (data not shown).

### Mechanical competition contributes to the competitive proficiency of cancer cells

To further characterize the phenotype of our cell competition model, we studied the orthogonal organization of cells competing in 2D culture. After various days of coculture, we stained cells with phalloidin and DAPI to visualize the cell membrane and the nucleus, respectively. Whereas AML12^EGFP^ cells in culture with AML12^TdT^ cells were organized as a monolayer for the whole duration of the experiment, they frequently formed bi-layered and tri-layered structures at the interface with cancer cells [Fig. S7]. This phenomenon may indicate that cancer cells exert compressive forces that dislocate and squeeze together AML12 cells, especially when cell crowding is more pronounced at day 9. To verify specifically this hypothesis, we conducted particle image velocimetry (PIV) to measure the compaction experienced by AML12^EGFP^ cells over time in movies acquired through time-lapse microscopy from day 3 to day 5 of culture. Most cancer cell lines induced compaction of AML12^EGFP^ cells which was modestly to significantly higher than the one in the culture with AML12^TdT^ cells [Fig. 5A-B, supplementary video 2 and 3], although there was not a clear positive correlation between the competitive proficiency and the intensity of this compaction. Nevertheless, the strongest competitor cancer cell lines B16 cells, among murine tumors, and Panc1, among human tumors, were also the ones that generated the most intense compaction in each experimental group.

**Figure 5:**
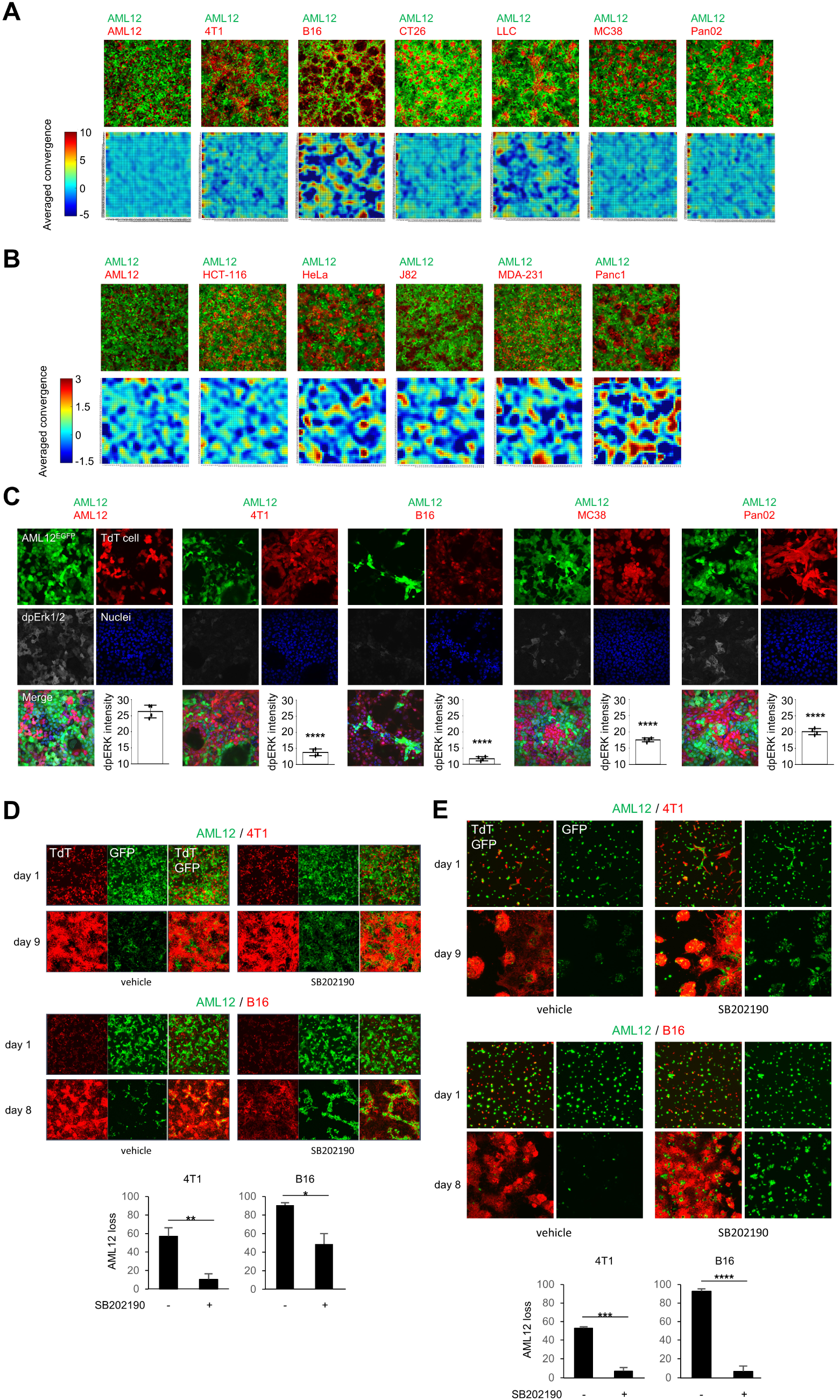
Study of mechanical cell competition. **(A)** Particle image velocimetry analysis of time-lapse microscopy acquisitions of AML12^EGFP^ cells in culture with murine cancer cells (day 3 to day 5). The upper panels show the first images of each series (time 0), while the heat maps show the average intensity of the convergence experienced by AML12^EGFP^ cells during the whole live imaging acquisition. **(B)** Particle image velocimetry analysis of AML12^EGFP^ cells in culture with human cancer cells (day 3 to day 5). **(C)** Representative images of cells stained with anti-phospho-ERK1/2 (Thr202/Tyr204) antibody. A chart in each panel shows the mean intensity of double phosphorylated ERK (dpERK) staining in AML12^EGFP^ cells. The values obtained in each culture with cancer cells are compared to the value from the culture with AML12^TdT^ cells. p-values for ANOVA test are shown. **(D)** Representative images of AML12^EGFP^ cells (green) in 2D culture with cancer cells expressing TdTomato (TdT, red) in media containing the p38 inhibitor SB202190 (10 μM) or vehicle. At the bottom, a chart shows the mean and the SEM of the loss of AML12^EGFP^ cells total area. p-values are obtained with unpaired T-test. **(E)** Representative images of AML12^EGFP^ cells (green) cultured in 3D with cancer cells (red) in media containing SB202190 (10 μM) or vehicle. At the bottom, a chart shows the mean and the SEM of the loss of the volume occupied by AML12^EGFP^ cells. p-values are obtained with unpaired T-test.

We recently reported that, in a *Drosophila melanogaster* model of mechanical cell competition, loser cells downregulate extracellular signal-regulated kinase 1/2 (ERK1/2) signaling before undergoing cell death [37]. Building upon this observation, we questioned whether ERK signaling is also impacted in AML12 cells cultured with cancer cells in 2D. After 4 days of culture, we immunostained cells to detect total ERK1/2 and its active form phosphorylated on Thr202 and Tyr204. The levels of total ERK in AML12^EGFP^ cells were reduced by approximately 30% when cultured with B16 cells compared to AML12^TdT^ cells, while there were no significant alterations in cultures with 4T1 cells, MC38 cells and Pan02 cells [Fig. S8A]. Double phosphorylated ERK (dpERK) levels exhibited a significant decrease in the cultures with cancer cells. Strong competitors B16 cells and 4T1 cells resulted in approximately 55% and 48% reduction, respectively, while poor competitors MC38 cells and Pan02 cells showed about 33% and 24% reduction, respectively [Fig. 5C].

Using the canine cell line MDCK as a model, it has been shown that compaction-driven cell competition activates the Rho-associated kinase ROCK1 and the stress kinase p38 in loser cells, which in turn leads to their elimination through upregulation of p53 [30]. Consistent with these findings, the pharmacological inhibition of p38 with SB202190 considerably prevented the elimination of AML12 cells when cultured with 4T1 cells and B16 cells, both in 2D culture [Fig. 5D] and to a greater extent in 3D culture [Fig. 5E]. This effect was not attributable to the inhibition of cancer cells growth, as their expansion in monoculture remained unaffected by SB202190 treatment [Fig. S8B]. Conversely, the pharmacological inhibition of other molecular processes, known to mediate the elimination of loser cells in various competitive scenarios, provided none to modest and inconsistent protection for AML12 cells in our competition assay. These interventions included the inhibition of entosis [22,24] using the ROCK inhibitor Y-27632 [Fig. S8C], caspase-dependent cell death [9,10] with the pan-caspase inhibitor Emricasan [Fig. S8D-E], and necroptosis [27] via the inhibition of receptor-interacting serine/threonine-protein kinase 1 (RIPK1) with Nec1s [Fig. S8D-E]. Finally, the mRNA expression level of the fitness fingerprint Fwe-Win isoforms, namely Fwe-2 and Fwe-4 [11], was not higher in cancer cells compared to AML12 cells in coculture [Fig. S8F], and the knock out (KO) of the human Fwe gene in HCT-116 cells did not protect AML12 cells from elimination [Fig. S8G]. In summary, these data collectively suggest that mechanical competition is a plausible mechanism enabling cancer cells to outcompete hepatocytes *in vitro*.

## Discussion

An established cancer represents the late phase of a series of events that often initiate many years earlier, and lead transformed cells to progressively acquire new traits and functions. Among these changes, the development of a supercompetitor state is considered crucial for the progression of the tumor [7,8,13,14]. In this research, our primary focus was to investigate the competitive interactions between cancer cells and hepatocytes. This choice was motivated by several reasons. First, the liver is a common metastatic site for numerous types of cancer [40], meaning that heterotypic cell competition between cancer cells and hepatocytes is a relevant phenomenon. Second, the liver is mostly constituted by solid parenchyma, which makes it an optimal organ for investigating tumor infiltrative growth. Third, hepatocytes represent the predominant cellular population in the liver, constituting up to 80% of its mass [29]. Therefore, we reasoned that exploring their competitive interaction with cancer cells could serve as a surrogate for understanding the dynamics within liver metastases.

Although cancer cells are generally considered to be intrinsically supercompetitors [5,14], our study of heterotypic cell competition reveals that the majority of cancer cell lines exhibit poor competitiveness towards hepatocytes *in vitro*. Importantly, we also found that the competitive strength of cancer cells can vary between 2D culture and 3D culture, possibly due to distinct competitive pressures experienced by cells in such different growth conditions. This finding has relevant implications in the field, since an amount of data describing the competition of mammalian cells *in vitro* have been derived only from cells cultured as monolayers [6,10,24,30–32].

Limited data are available describing heterotypic cell competition in mammalian organisms. Moya et al recently discovered that primary liver tumors compete with neighboring hepatocytes based on the activation level of YAP and TAZ, the downstream effectors of the Hippo pathway [35]. Interestingly, increasing the activation of YAP in hepatocytes after the settlement of metastatic melanoma cells resulted in a remarkable 98% decrease in liver metastases load compared to mice with unperturbed YAP activation. Recently, our group reported that inhibiting fitness fingerprint-mediated cell competition, via stable knockdown of Fwe expression in human cancer cell lines, significantly reduced the incidence of spontaneous metastases to a number of organs following the subcutaneous implantation of tumor cells [10]. Our new data now suggest that poor competitor cancer cells which do not eliminate hepatocytes *in vitro* can turn into supercompetitors *in vivo* and replace the liver parenchyma. Moreover, there might be a correlation between the competitive competence of cancer cells and the behavior of liver metastases in terms of aggressiveness and morphology. In fact, the two strong competitor cancer cell lines in our panel generated aggressive metastases with an infiltrative morphology, whereas most of the poor competitor cancer cell lines resulted in a more indolent disease which often displayed expansile growth. Further studies are needed to validate this observation. Intriguingly, a number of reports consistently found that liver metastases with infiltrative/replacing growth pattern are associated with worse prognosis compared to those with expansive or desmoplastic growth in patients with colorectal cancer [43], pancreatic cancer [44] and cutaneous melanoma [45]. This aligns with our findings in mouse models, and suggests that preventing the elimination of healthy cells might alter the growth pattern of various aggressive tumors and hinder their progression.

Increased homeostatic pressure is thought to be a characteristic trait of solid tumors, due to their uncontrolled growth within tissues that are subjected to space constraints [23,39]. Mechanical cell competition occurs when clones of cells overproliferate, causing tissue crowding. This, in turn, generates mechanical forces across the tissue, particularly at the interface between cell populations with different growth rates. The different level of resilience to these mechanical forces results in the elimination of cells that are more sensitive to mechanical stress [36]. Mechanical competition has been well characterized in *Drosophila melanogaster* during development, where the measurement of tissue compaction with PIV together with the quantification of ERK signaling activation in cells have proven valuable approaches for its study [9,37]. In the present work, both PIV analysis and the assessment of ERK activation levels corroborate the existence of varied degrees of mechanical compression experienced by AML12 cells in cultures with cancer cells. Notably, the most formidable competitor within our panel, B16 cells, elicited the highest compaction intensity and the most significant reduction in ERK activity in AML12 cells. This suggests that mechanical competition may play a particularly prominent role in this specific tumor model.

At last, our data show that the inhibition of the kinase activity of p38 was sufficient to rescue the elimination of AML12 cells in culture with strong competitor cancer cells. Importantly, the fact that cancer cells still expanded without eliminating AML12 cells in the presence of a p38 inhibitor suggests that the outcompetition of healthy cells is not inherently coupled to the enlargement of the population of tumor cells. The main physiological function of p38 kinases is mediating cell survival in response to stress at many levels, including both environmental and intracellular stresses [41]. However, sustained activation of p38 has been shown to be detrimental for the survival of hepatocytes under certain stress conditions [42]. This opens the possibility that, besides protecting from mechanical competition, the inhibition of p38 in our system might prevent the elimination of AML12 cells by reducing the stress response resulting from the culture with cancer cells. This compelling hypothesis warrants further investigation.

In conclusion, this work represents a pivotal advance in the study of tumor heterotypic cell competition with healthy cells. Our study not only fills a critical gap in our understanding of cancer behavior but also opens new avenues for translational research and therapeutic exploration. The quest for identifying the molecular intricacies leading to the elimination of healthy cells and their replacement with cancer represents a promising frontier in oncological research, with the potential to revolutionize our approach to cancer therapeutics.

## Material and Methods

### Cell lines

AML12 cells (#CRL-2254), 4T1 cells (#CRL-2539), LLC cells (#CRL-1642) and Renca cells (#CRL-2947) were purchased from ATCC, while MC38 cells (#ENH204) were purchased from Kerafast. HCT-116^FweKO^ cells were previously developed in our lab [10]. The remaining cell lines were a gift from other groups at the Champalimaud Centre for the Unknown: B16 F10 cells, CT26 cells, EO771 cells, Pan02 cells, HCT-116 cells, Panc1 cells and SK-MEL-28 cells from Bruno Costa-Silva, MDA-231 cells from Rita Fior, Renca cells from Henrique Veiga-Fernandes, and HeLa cells, J82 cells and 293T cells from Mireia Castillo-Martin. Murine cancer cell lines were authenticated through STR profiling (ATCC #137-XV). All cells were regularly tested for Mycoplasma contamination.

AML12 cells were grown in DMEM F12 supplemented with fetal bovine serum (FBS) 10%, insulin-transferrin-selenium 1X (Corning #25-800-CR) and Dexamethasone 40 ng/mL. This media was also used for the coculture of AML12 cells with cancer cells. Cancer cells B16 F10, EO771, LLC, MC38, HCT-116, MDA-231 and Panc1 were grown in DMEM supplemented with 10% FBS, while 4T1, CT26, Pan02 and Renca in RPMI with 10% FBS, and HeLa, J82 and SK-MEL-28 in EMEM with 10% FBS.

### Drugs

SB202190 was purchased from Santa Cruz (#sc-202334), Emricasan (#S7775) and Y-27632 (#S1049) were purchased from Selleckchem, Nec1s was purchased from Biovision (#2263). All drugs were diluted in DMSO and were added to the growth media starting 1 day after cell seeding.

### Generation of cells expressing EGFP or TdTomato

293T cells were transfected using lipofectamine 3000 (Invitrogen #L3000015) with ViraPower packaging vectors (Invitrogen) and viral backbones. The supernatant was collected after 48 hours, filtered through 0.45 µm, and directly used to transduce AML12 cells and cancer cells. Clones expressing EGFP (FUGW virus; Addgene #14883) and TdTomato (FUtdTW virus; Addgene #22478) were purified by fluorescence-activated cell sorting (FACS) at the flow cytometer.

### Cell competition assay in 2D

AML12^EGFP^ cells (1.8×10^5^ cells per well) were seeded with murine cancer cells expressing TdT (6×10^4^ cells per well) in a 24-well µ-plate (Ibidi #82406). The growth media was replaced every day with fresh media. Human cancer cells were seeded 9×10^4^ cells per well.

### Cell competition assay in 3D

AML12^EGFP^ cells (3×10^4^ cells per well) were seeded with murine cancer cells expressing TdT (1.5×10^4^ cells per well) on top of growth-factor-reduced Matrigel (Corning #356252; 160 µL per well) in a 24-well µ-plate, and covered with growth media containing 5% Matrigel. Starting from day 2, the media was replaced every day with complete media. The number of cells seeded was increased by 50% in the coculture of AML12 cells with human cancer cells.

For the fully embedded growth condition, 80 µL of Matrigel per well were allowed to solidify for 30 minutes inside the incubator, then AML12^EGFP^ cells (3×10^5^) and cancer cells expressing TdT (1×10^5^) were resuspended in 220 μL of Matrigel and seeded. After 45 minutes, 1 mL of complete media was added on top, and was replaced every day. For imaging, a z-stack of approximately 400 µm was acquired.

### Proliferation analysis

Cells were stained with Cell Trace Violet (CTV; Invitrogen #C34557) following the manufacturer protocol for cells in suspension, then seeded in 2D in a 24-well plate either alone (4×10^4^ cells per well) or in coculture with confluent AML12^EGFP^ cells (cancer cells 6×10^4^ per well, AML12^EGFP^ cells 2.2×10^5^ per well). The flow cytometry data were analyzed with FlowJo v10.8.1.

### Crystal violet assay

Cells were seeded in a 96-well plate in quadruplicate (2×10^3^ cells per well). After 72 hours, cells were washed with PBS, incubated with 50 µL of crystal violet solution 0.5% (Fisher #11435027) for 20 minutes, washed in tap water and left to dry overnight. Methanol was added 200 µL per well, followed by incubation on an orbital shaker for 20 minutes. Absorbance was measured at 570 nm with a spectrophotometer.

### Treatment with conditioned medium

The conditioned medium was generated by growing confluent cells in poor DMEM F12 for 24 hours. Supernatant was filtered through 0.45 µm and mixed with the complete growth media of AML12 cells to generate a 30% solution, which was added to AML12^EGFP^ cells monocultures the day after seeding and was replaced every 2 days.

### Treatment with the supernatant from competing cells

AML12^EGFP^ cells were seeded in culture with cancer cells or AML12^TdT^ cells in 2D. The media was replaced after 2 days with fresh media, and after 24 hours the supernatant was collected and filtered through 0.45 µm. It was then mixed with the complete growth media of AML12 cells to generate a 30% solution, which was added to AML12^EGFP^ cells monocultures the day after seeding and was replaced every 2 days.

### Transwell culture

AML12^EGFP^ cells (2.4×10^5^) were seeded on top of inserts with a pore diameter of 0.4 μm (Thincert #665640) which were placed inside a 12-well plate containing at the bottom cancer cells expressing TdT (1.2×10^5^ cells per well). The growth media was replaced every day.

### Annexin V assay

Cells were collected, washed, and stained for 15 minutes with Annexin V Pacific Blue (Biolegend #640918) 2 µL/100 µL and propidium iodide (Sigma #P4170) 20 µg/mL in Annexin V Binding Buffer (Biolegend #422201). Cells were then run into a flow cytometer and the results were analyzed using FlowJo. To identify cell debris, we displayed the Annexin V channel and the PI channel, and selected the double negative population. This was plotted according to the forward scatter (FSC) and the side scatter (SSC), and the events in the region with low FSC, which include cell debris, were excluded (ungated). The resulting population was selected for single cells, then AML12^EGFP^ cells were gated and analyzed for positivity to Annexin V and PI. Experiments were conducted in triplicate and repeated three times.

### Phalloidin staining

AML12^EGFP^ cells 3.3×10^4^ and cells expressing TdT 1.1×10^4^ were seeded in 18-well glass bottom µ-slides (Ibidi #81817). After 1 day, 5 days and 9 days, cells were fixed with PFA 4% in PBS for 15 minutes at room temperature, permeabilized with Triton X-100 0.1% for 15 minutes, and stained for 45 minutes with a solution of PBS 1% bovine serum albumin (BSA) containing Phalloidin Alexa Fluor Plus 647 (Invitrogen #A30107) 1X and DAPI (Thermofisher #62248) 1 µg/mL.

### Immunofluorescence

Cells were cultured in 18-well glass bottom µ-slides (Ibidi #81817). After 4 days, cells were fixed with PFA 4% in PBS for 15 minutes at room temperature, permeabilized with Triton X-100 0.1% for 15 minutes, blocked for 60 minutes in PBS 1% BSA, and stained with anti-Erk1/2 antibody (CST #4695) 1:800 or anti-phospho-Erk1/2 Thr202/Tyr204 antibody (CST #4370) 1:300, both diluted in PBS 1% BSA. After 60 minutes of incubation at room temperature, cells were washed with PBS 0.05% Tween (PBST), then incubated for 60 minutes with the secondary antibody anti-Rabbit 647 (Invitrogen #A-31573) 1:1000 in PBS 1% BSA. After washing with PBST, cells were incubated for 5 minutes in PBS containing DAPI 1 µg/mL.

### RT-PCR

Cells in coculture in 2D for 3 days were collected, sorted via FACS, and total RNA was extracted using RNeasy Mini Kit (Qiagen #74104). One µg of RNA was used to generate cDNA with QuantiTect Reverse Transcription kit (Qiagen #205313). For the RT-PCR reaction, we used PowerUp SYBR Green Master Mix (Applied Biosystems #A25777) with the following primers:

mFwe1 Fw ATCTGTCGGCCAAGCTAACC; Rv GGGAAGTAACTGAGTCGCGT

mFwe2 Fw GGAACTGTGAGGCCTGGAG; Rv GCCAGTGAAACAGCTCTCCT

mFwe3 Fw CTGTCTTCTACTGCGGGCAT; Rv TTTCTGTTCGGCAGTCTCACA

mFwe4 Fw TGCTAAATCCTGGGTGTCCC; Rv GAGGGTGGATAGTGACGCAG

18S Fw GTAACCCGTTGAACCCCATT; Rv CCATCCAATCGGTAGTAGCG

### Experimental liver metastases and histopathology

Murine cancer cells were diluted in PBS and 1×10^6^ cells in 100 µL were injected into the spleen of 9-10 weeks old syngeneic mice (n. 4 per cell line; C57BL/6 mouse for B16 F10 cells, EO771 cells, LLC cells, MC38 cells and Pan02 cells; BALB/c mouse for 4T1 cells, CT26 cells and Renca cells), followed by splenectomy. Mice were sacrificed after 21 days, or earlier when the first 2 mice of the experimental group reached humane endpoints. For tumor xenografts, 1×10^6^ human cancer cells (HCT-116, HeLa, MDA-231, Panc1 and SK-MEL-28) were diluted in PBS and injected into the spleen of nude mice (n. 3 per cell line), which were sacrificed after 28 days.

Liver was formalin-fixed, paraffin-embedded, sectioned (4 μm) and stained with hematoxylin and eosin. Sections were analyzed by a pathologist blinded to the experimental group in an Axioscope 5 microscope (Zeiss), and scored for the criteria shown in Table 1, adapted from Friedl et al [38]. Briefly, pattern of tumor growth, tumor border configuration, intercalation between tumor cells and hepatocytes, and the presence of liver damage were annotated for all mice in all experimental groups. Microphotographs were captured with an Axiocam 208 color camera (Zeiss).

Quantification of hepatocyte and tumor cell apoptosis was performed in liver slides by anti-cleaved caspase-3 immunostaining (Cell Signaling Technology #9661, diluted 1:400), detected with HRP-bound goat anti-rabbit secondary antibody (Invitrogen #31460) followed by DAB substrate kit (VectorLabs #SK-4100), and counterstained with hematoxylin.

### Microscopy imaging and analysis

Cell culture images were acquired with a Zeiss LSM 880 confocal microscope. To excite the GFP and TdTomato fluorophores, the Argon generated 488 nm and the DPSS 561 nm laser lines were utilized, respectively. Images were acquired using a 10x/0.45 objective lens (unless otherwise specified) and the signal was collected by using a highly efficient GaAsP detector. The quantification of the area of fluorescent cells in 2D culture was done with Fiji, and is based on the maximum intensity projection and manual segmentation. The quantification of the volume of cells in 3D culture was done with Imaris v9.3.1 using manual segmentation.

Particle image velocimetry (PIV) was calculated using a previously developed PIVlab MATLAB routine [46]. To obtain higher accuracy in the displacement, two passes analysis was applied. For the first pass the interrogation window was set to 128 pixels with 50%, and for the second the window size was set to 64 pixels with 50% overlap. Parameters for this PIV were analysis boxes of 64 pixels (∼44×44 mm), 50% overlap boxes. PIV displacements at time t were determined based on images t-25 and t+25min. Displacement maps were spatially filtered to remove noise coming from outliers. The convergence was calculated as the opposite of the field divergence and the average value over all frames is shown in the results.

The quantification of the intensity of the staining for Erk1/2 and dpErk1/2 in AML12^EGFP^ cells was done using a custom Fiji macro. The pipeline in the macro calculates a binary mask of AML12^EGFP^ cells by normalizing their intensity and applying a triangle thresholding method. These masks are used to calculate the staining for Erk1/2 and dpErk1/2 on AML12^EGFP^ cells.

### Statistical analysis

For all the experiments, normal distribution was assessed using the Shapiro-Wilk test. Two-tailed unpaired Student T-test was used for comparison of two groups, while ANOVA test was chosen for comparison of multiple groups.

## Acknowledgements

We thank Bruno Costa-Silva, Mireia Castillo-Martin, Henrique Veiga-Fernandes and Rita Fior for providing various cancer cell lines used in this study. We also acknowledge Anna Pezzarossa for the technical assistance in the imaging analysis. We thank Catarina Brás-Pereira for her invaluable support and critical feedback during the execution of this research. This study has been funded by the European Research Council (Consolidator Grant to E.M.: ‘‘Active Mechanisms of Cell Selection: From Cell Competition to Cell Fitness’’) and the Champalimaud Foundation.

## Author contributions

A.S. and E.M. conceptualized the idea, designed and interpreted the experiments. A.S. performed and analyzed the experiments, and wrote the manuscript. T.C. provided the pathological description of liver metastases. M.A.F.P. analyzed the data shown in Fig. 5B. M.M.R. generated the Fig. 4D and, together with A.G.G., edited the Supplementary videos. D.A. helped in the acquisition and analysis of images.

