## Supplementary Figures for "Heterotypic competition between cancer cells and hepatocytes generates heterogeneous context-dependent phenotypes"

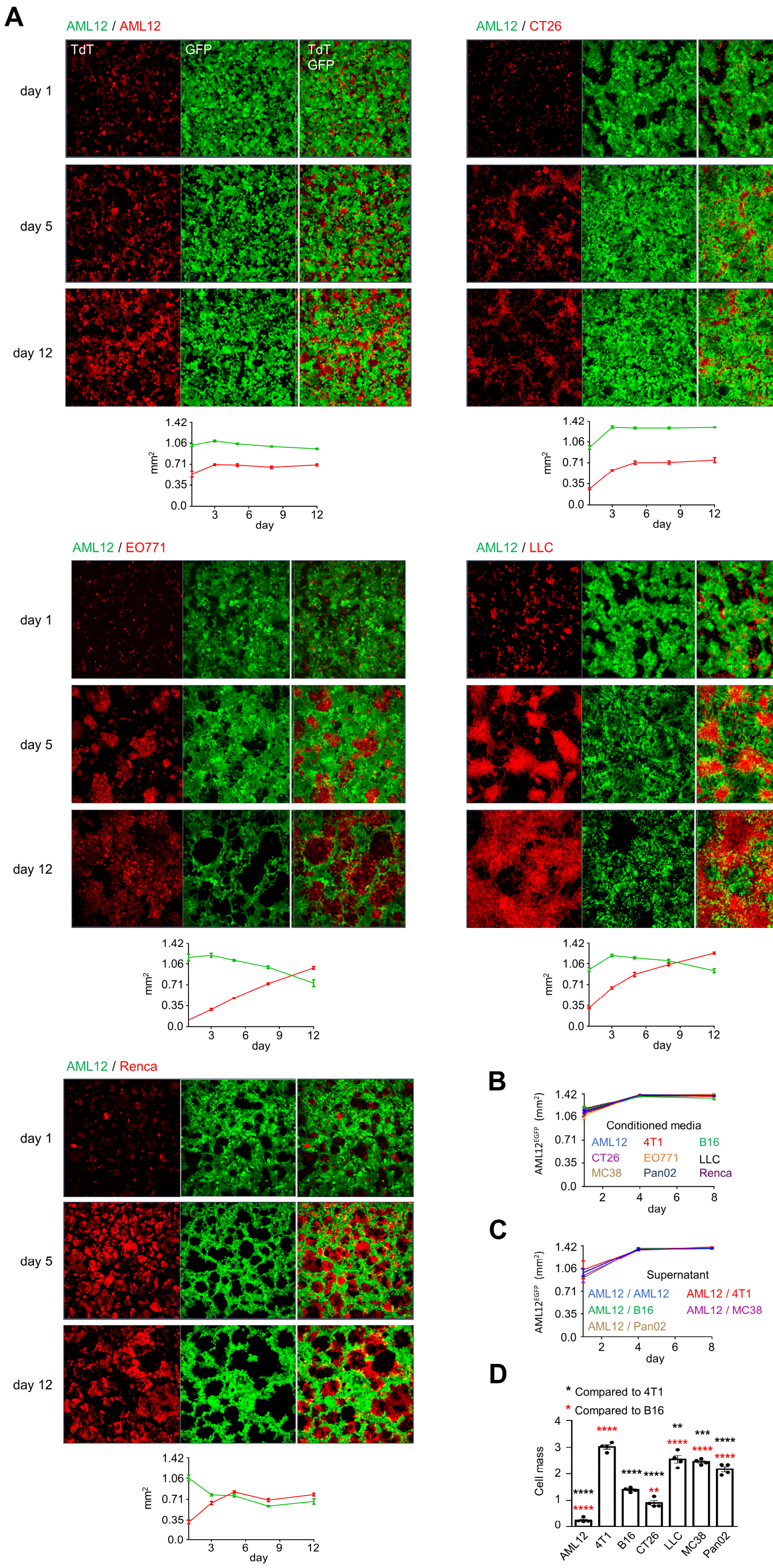

A
AML12 / AML12
TdT
GFP
TdT GFP
day 1
day 5
day 12
AML12 / CT26
AML12 / EO771
AML12 / LLC
AML12 / Renca
mm2
day
B
AML12EGFP (mm2)
Conditioned media
AML12
4T1
B16
CT26
EO771
LLC
MC38
Pan02
Renca
C
Supernatant
AML12 / AML12
AML12 / 4T1
AML12 / B16
AML12 / MC38
AML12 / Pan02
D
* Compared to 4T1
* Compared to B16
Cell mass
AML12
4T1
B16
CT26
LLC
MC38
Pan02

**Supplementary figure 1. (A)** Representative images, shown as maximum intensity projection, of AML12$^{EGFP}$ cells (green) in 2D culture with murine cancer cells expressing TdTomato (TdT, red). At the bottom of each panel, a chart shows the area per field of view occupied over time by AML12$^{EGFP}$ cells (green curve) and the partner cell line (red curve). Each dot shows the mean and the SEM of the values from at least 4 images. **(B)** Total area per field of view occupied by AML12$^{EGFP}$ cells during treatment with the conditioned media derived from cancer cells. **(C)** Total area per field of view occupied by AML12$^{EGFP}$ cells during treatment with the supernatant from the indicated cocultures. **(D)** Cell mass quantification by spectrophotometry of cells grown in monoculture for 72 hours and stained with Crystal Violet. p-values calculated with ANOVA test.

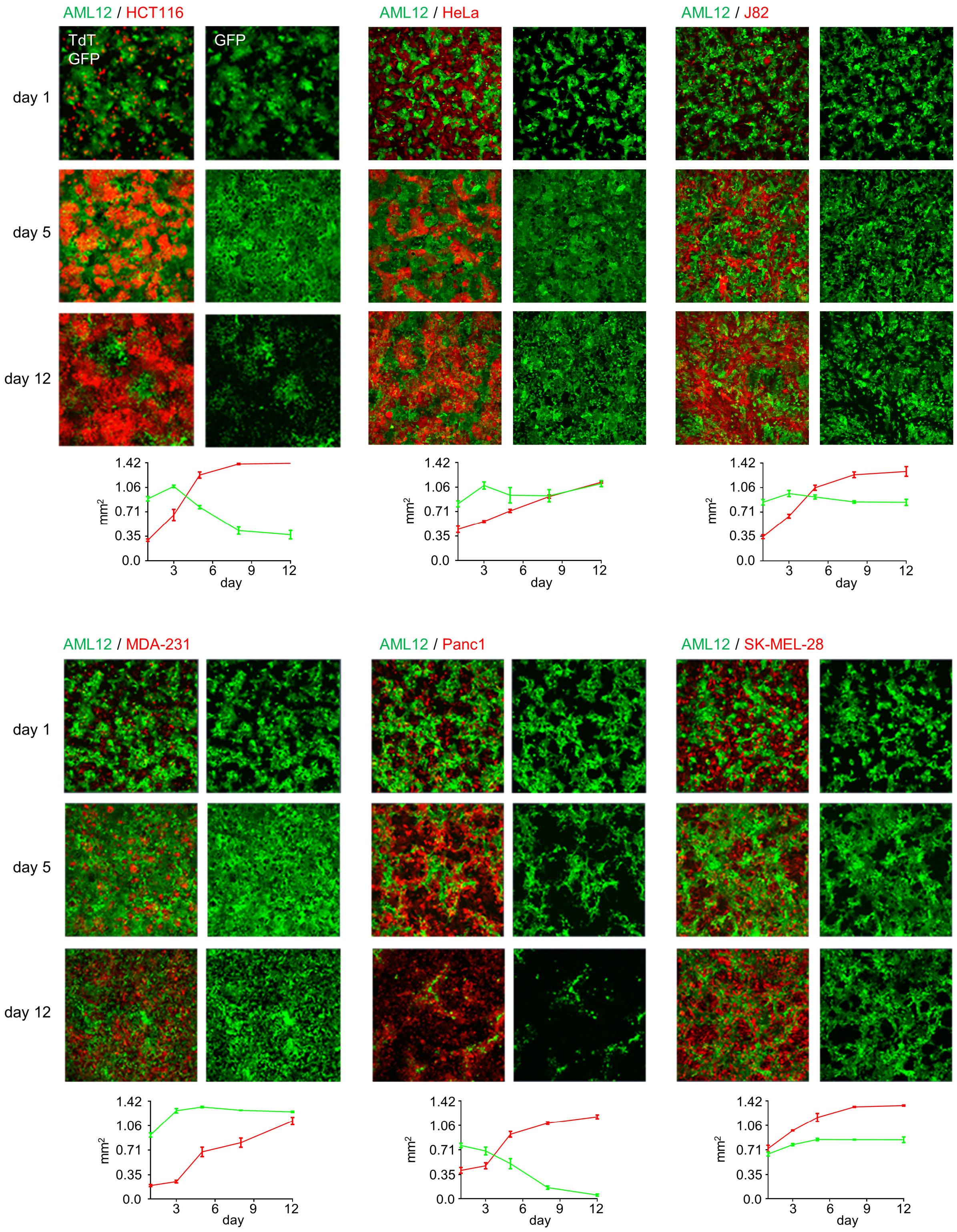
AML12 / HCT116
TdT
GFP
GFP
AML12 / HeLa
AML12 / J82
day 1
day 5
day 12
1.42
1.06
0.71
0.35
0.0
mm$^2$
3
6
9
12
day
AML12 / MDA-231
AML12 / Panc1
AML12 / SK-MEL-28

**Supplementary figure 2.** Representative images, shown as maximum intensity projection, of AML12$^{EGFP}$ cells (green) in 2D culture with human cancer cells expressing TdTomato (TdT, red). At the bottom of each panel, a chart shows the area per field of view occupied over time by AML12$^{EGFP}$ cells (green curve) and the partner cell line (red curve). Each dot shows the mean and the SEM of the values from at least 3 images.

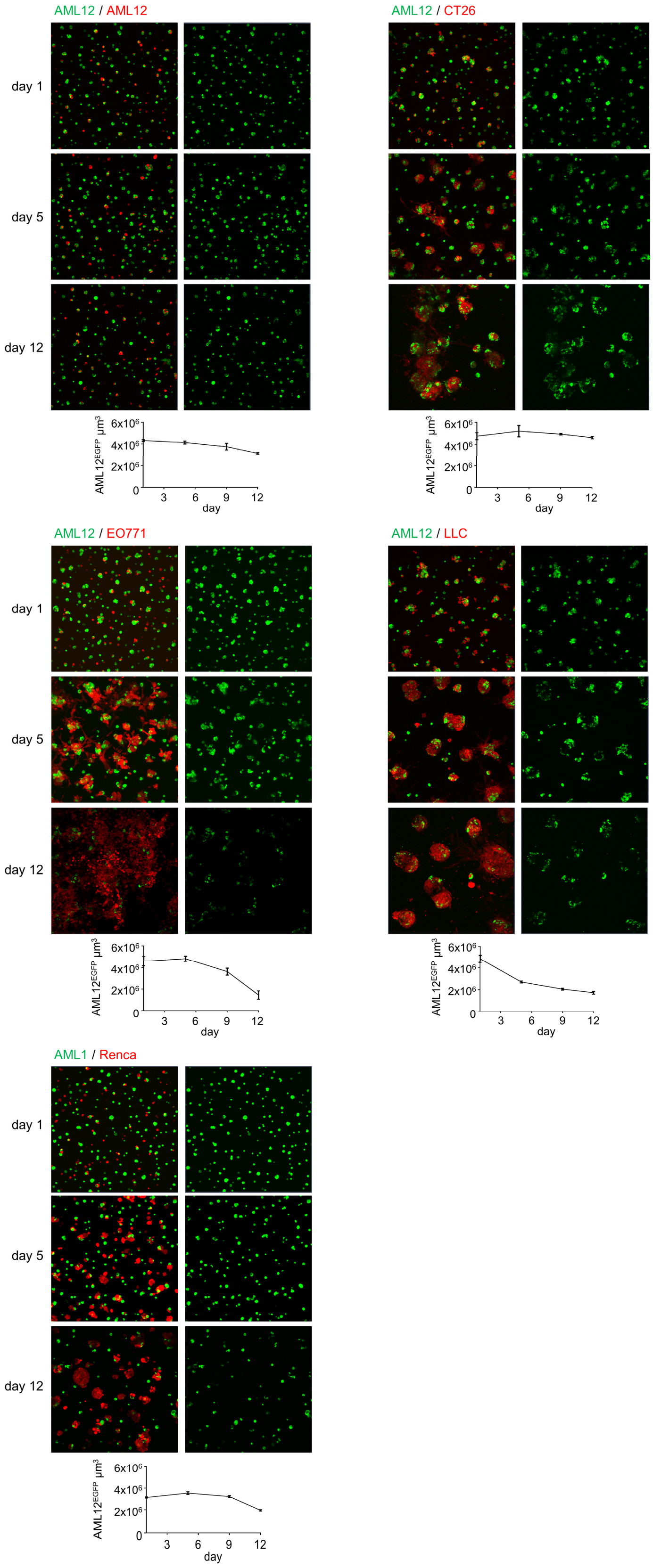
AML12 / AML12
day 1
day 5
day 12
AML12EGFP µm3
6x106
4x106
2x106
0
3
6
9
12
day
AML12 / CT26
AML12 / EO771
AML12 / LLC
AML1 / Renca

**Supplementary figure 3.** Representative images, shown as maximum intensity projection, of AML12$^{EGFP}$ cells (green) in culture on top of Matrigel gel with murine cancer cells marked with TdTomato (TdT, red). At the bottom of each panel, a chart shows the total volume per field of view (10x objective) occupied over time by AML12$^{EGFP}$ cells. Each dot shows the mean and the SEM of the values from at least 3 images.

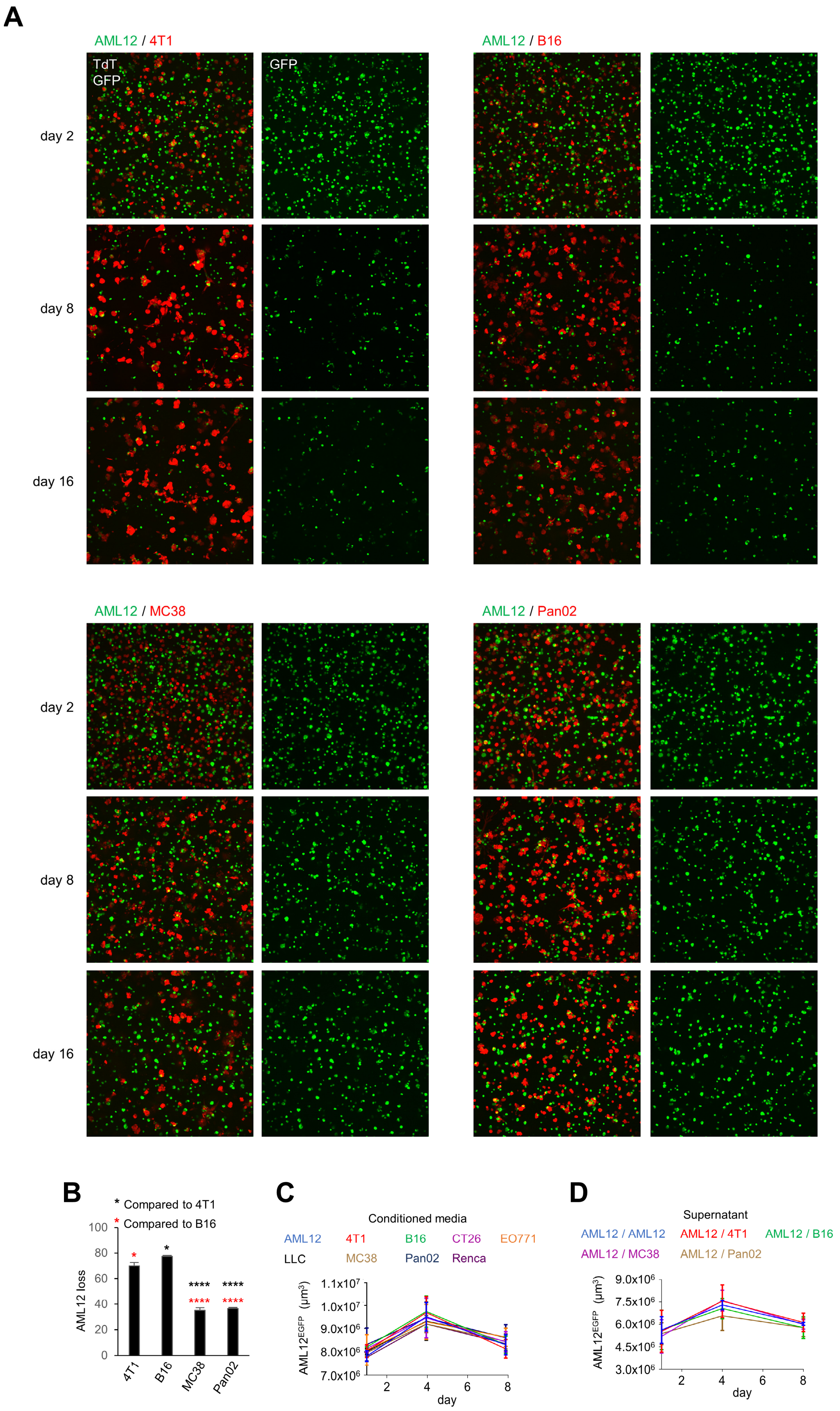
A
AML12 / 4T1
TdT
GFP
GFP
AML12 / B16
day 2
day 8
day 16
AML12 / MC38
AML12 / Pan02
day 2
day 8
day 16
B
* Compared to 4T1
* Compared to B16
AML12 loss
4T1
B16
MC38
Pan02
C
Conditioned media
AML12
4T1
B16
CT26
EO771
LLC
MC38
Pan02
Renca
AML12EGFP (µm3)
day
D
Supernatant
AML12 / AML12
AML12 / 4T1
AML12 / B16
AML12 / MC38
AML12 / Pan02
AML12EGFP (µm3)
day

**Supplementary figure 4. (A)** Representative images, shown as maximum intensity projection of AML12$^{EGFP}$ cells (green) fully embedded in Matrigel gel with cancer cells marked with TdT (red) and visualized with a 10x objective across a Z-stack of 400 µm. **(B)** Quantification of the loss of AML12$^{EGFP}$ cells from the experiment in A, measured as the ratio $\frac{Volume\ AML12\ day\ 16}{Volume\ AML12\ day\ 2}$. p-values obtained with ANOVA. **(C)** Total volume per field of view occupied by AML12$^{EGFP}$ cells during treatment with the conditioned media derived from cancer cells. **(D)** Total volume per field of view occupied by AML12$^{EGFP}$ cells during treatment with the supernatant from the cocultures indicated on the top.

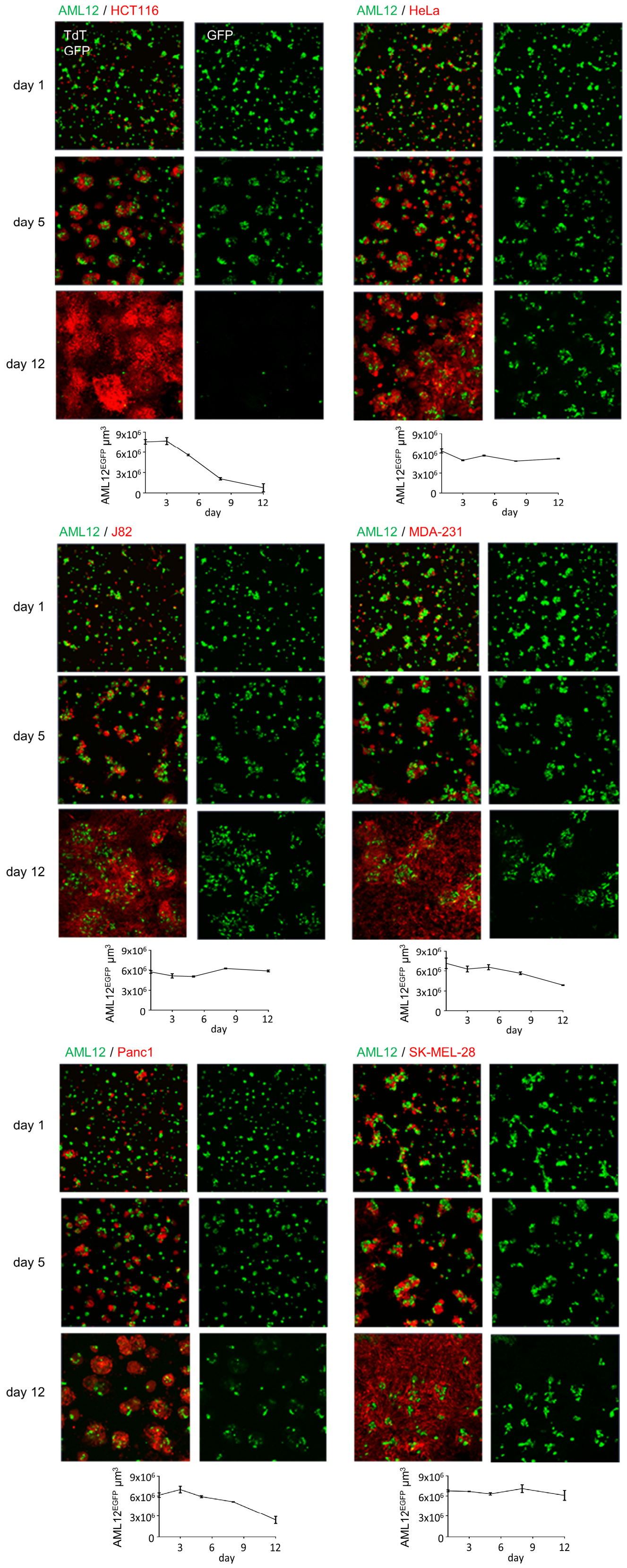
AML12 / HCT116
TdT
GFP
GFP
day 1
day 5
day 12
AML12EGFP µm3
9x106
6x106
3x106
0
3
6
9
12
day
AML12 / HeLa
AML12 / J82
AML12 / MDA-231
AML12 / Panc1
AML12 / SK-MEL-28

**Supplementary figure 5.** Representative images, shown as maximum intensity projection, of AML12$^{EGFP}$ cells (green) in culture on top of Matrigel gel with human cancer cells marked with TdTomato (TdT, red). At the bottom of each panel**,** a chart shows the volume per field of view occupied over time by AML12$^{EGFP}$ cells. Each dot represents the mean and the SEM derived from at least 3 different photos.

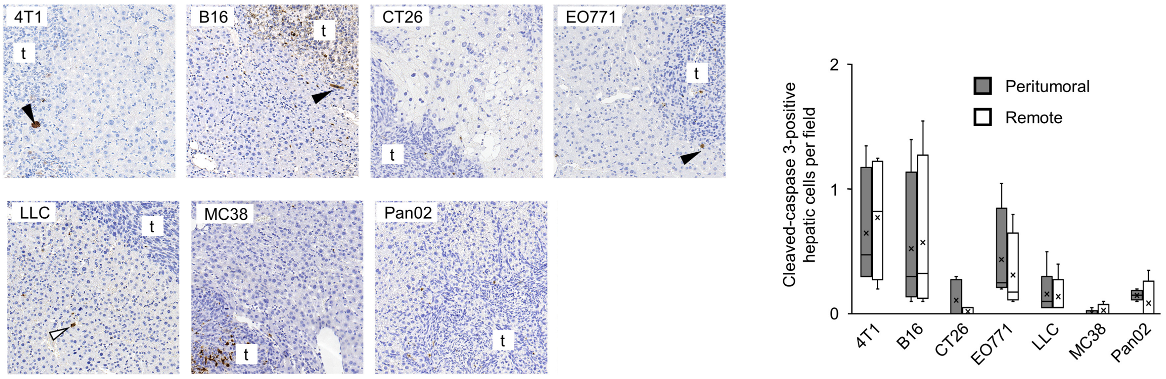
4T1
t
B16
t
CT26
t
EO771
t
LLC
t
MC38
t
Pan02
t
Cleaved-caspase 3-positive hepatic cells per field
Peritumoral
Remote
2
1
0
4T1
B16
CT26
EO771
LLC
MC38
Pan02

**Supplementary figure 6.** Representative images of liver sections stained with an anti-cleaved caspase-3 (cCasp-3) antibody and counterstained with hematoxylin. The plot shows the number of cCasp-3 positive hepatic cells at the tumor border and far from the tumor per field of view. Black arrowhead: peritumoral cCasp-3-positive hepatocytes. White arrowhead: remote cCasp-3-positive cells.

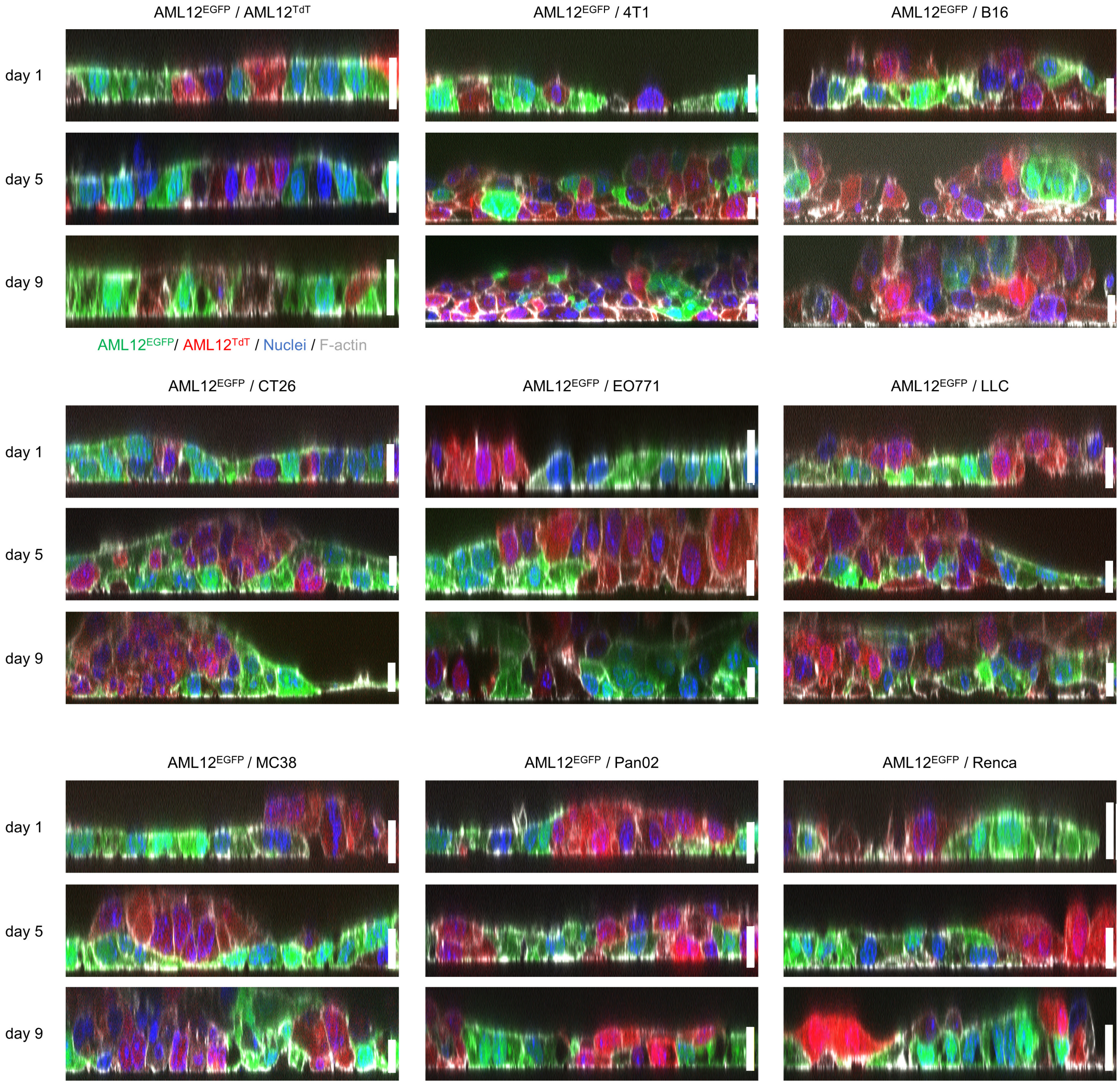
AML12EGFP / AML12TdT
AML12EGFP / 4T1
AML12EGFP / B16
day 1
day 5
day 9
AML12EGFP/ AML12TdT / Nuclei / F-actin
AML12EGFP / CT26
AML12EGFP / EO771
AML12EGFP / LLC
AML12EGFP / MC38
AML12EGFP / Pan02
AML12EGFP / Renca

**Supplementary figure 7.** Representative images of the XZ plane of AML12$^{EGFP}$ cells (green) in culture with murine cancer cells expressing TdTomato (TdT, red). Images were acquired with a 63x objective (scale bar 10 μm) after staining with phalloidin (grey), to visualize the cell membrane, and DAPI (blue), to mark the nucleus.

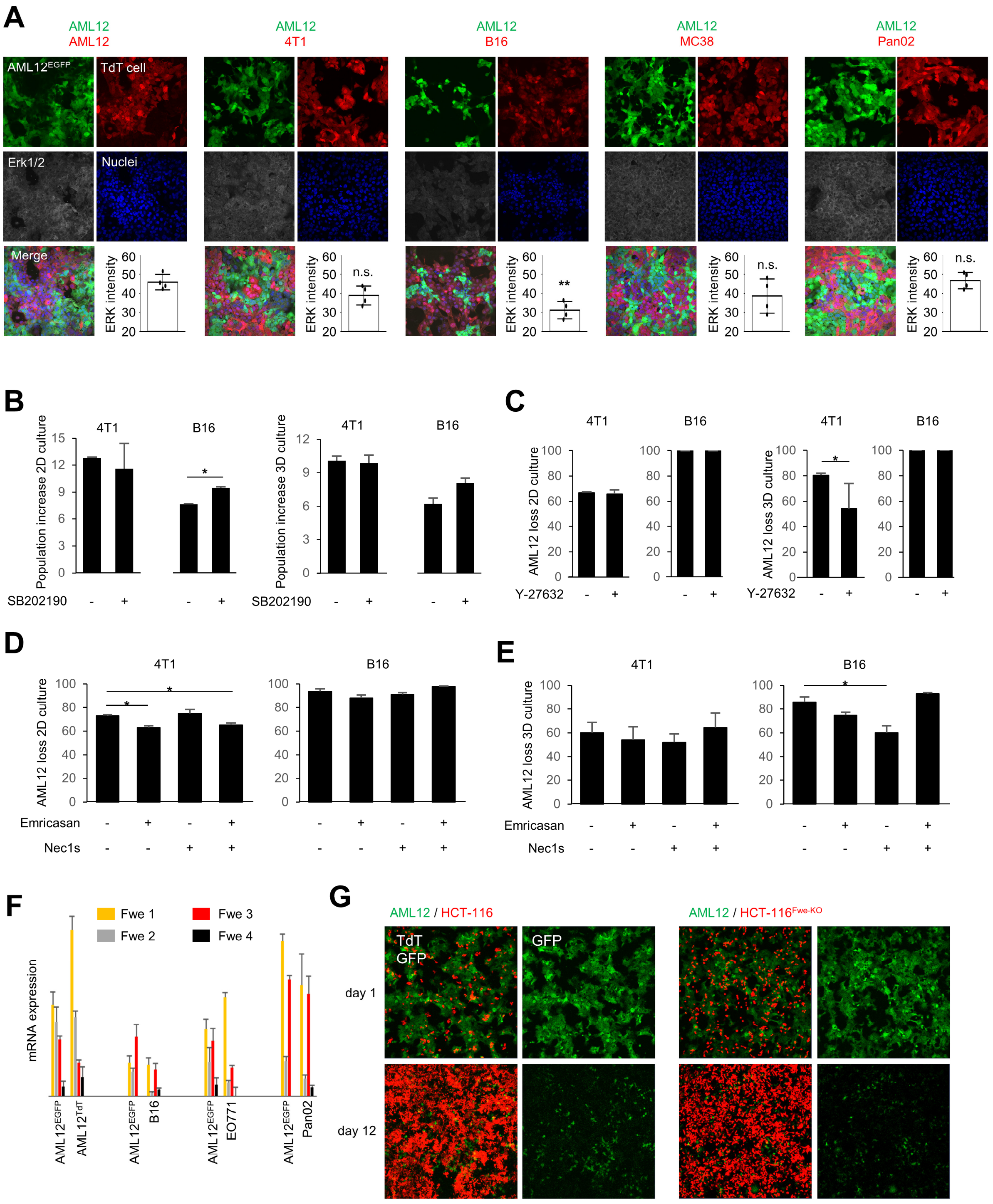
A
AML12
AML12
4T1
B16
MC38
Pan02
AML12EGFP
TdT cell
Erk1/2
Nuclei
Merge
ERK intensity
n.s.
**
B
Population increase 2D culture
Population increase 3D culture
4T1
B16
SB202190
C
AML12 loss 2D culture
AML12 loss 3D culture
Y-27632
D
AML12 loss 2D culture
Emricasan
Nec1s
E
AML12 loss 3D culture
F
Fwe 1
Fwe 2
Fwe 3
Fwe 4
mRNA expression
AML12EGFP
AML12TdT
B16
EO771
Pan02
G
AML12 / HCT-116
AML12 / HCT-116Fwe-KO
TdT
GFP
day 1
day 12

**Supplementary figure 8. (A)** Representative images of cells stained with an anti-ERK1/2 antibody. A chart in each panel shows the mean intensity of ERK1/2 staining in AML12$^{EGFP}$ cells. The values obtained in the cultures with cancer cells are compared to that from the culture with AML12$^{TdT}$ cells. p-values obtained with ANOVA. **(B)** Quantification of the total area per field of view (left panel, 2D culture of 5 days) and of the total volume per field of view (right panel, 3D culture of 8 days) of 4T1 cells and B16 cells monocultures treated with SB202190 (10 μM) or vehicle. **(C)** Quantification of the loss of the total area per field of view (left panel, 2D culture of 12 days) and of the total volume per field of view (right panel, 3D culture of 12 days) of AML12$^{EGFP}$ cells in coculture with 4T1 cells and B16 cells and treated with Y-27632 (10 μM) or vehicle. **(D** and **E)** Quantification of the loss of the total area per field of view (C, 2D culture) and the total volume per field of view (D, 3D culture) of AML12$^{EGFP}$ cells in coculture for 10 days with B16 cells and 4T1 cells, in the presence of Emricasan (10 μM), Nec1s (20 μM), Emricasan + Nec1s, or vehicle. For each chart, the values are compared to those of control cells treated with vehicle. **(F)** mRNA expression levels of the 4 mouse Flower (Fwe) isoforms by real-time PCR (RT-PCR) in pair of cells cultured together for 3 days as monolayers. The y axis indicates the expression levels of each isoform after normalization for the expression of 18S. **(G)** Representative images of AML12$^{EGFP}$ cells in culture with HCT-116 cells wild type and HCT-116 cells that are knock out (KO) for the Fwe gene.

**Supplementary video 1:** Time-lapse sequence (lapse interval of 30 min) of $AML12^{EGFP}$ cells in 3D culture with murine cancer cells. Images show the maximum intensity projection.

**Supplementary video 2:** Time-lapse sequence (lapse interval of 30 min) of $AML12^{EGFP}$ cells in 2D culture with murine cancer cells.

**Supplementary video 3:** Time-lapse imaging (lapse interval of 20 min) of $AML12^{EGFP}$ cells in 2D culture with human cancer cells.
